# Carboxylated graphene: A novel approach for enhanced IgA-SARS-CoV-2 electrochemical biosensing

**DOI:** 10.1101/2024.02.27.582131

**Authors:** Luciana de Souza Freire, Camila Macena Ruzo, Barbara Batista Salgado, Ariamna María Dip Gandarilla, Yonny Romaguera Barcelay, Ana P.M. Tavares, Francisco Xavier Nobre, Pritesh Lalwani, Spartaco Astofi-Filho, Walter Ricardo Brito

## Abstract

Biosensors comprise devices that use a material of biological nature as receptors connected to transducers, these devices are capable of capturing biorecognition signals, called a primary signal, and converting it to a measurable signal. In this study, we report the synthesis of carboxylated graphene (CG) through a carboxylation method in acid medium and further characterization of the materials by different techniques such as scanning electron microscopy (SEM), energy-dispersive X-ray spectroscopy (EDS), Raman spectroscopy, thermal gravimetric analysis (TGA), and X-ray diffraction (DRX). Also, the surface of the screen-printed carbon electrodes (SPCEs) was modified with CG for subsequent immobilization of N-protein of SARS-CoV-2, which allowed the detection of antibodies (IgA-SARS-CoV-2). The electrical properties and response of the biosensor were investigated using electrochemical techniques (cyclic voltammetry and electrochemical impedance spectroscopy). Through the chemical characterization techniques, it was possible to confirm the success of the CG synthesis process. The biosensor fabricated shown to be able to detect IgA-SARS-CoV-2 in the range of 1:1000 to1:200 v/v in phosphate buffer solution (PBS) and the limit of detection calculated was 1:1601 v/v. this perspective they comprise a wide range of applications due to its advantages, such as the possibility of a shorter response time, reproducibility, the miniaturization of detection devices such as the use of screen-printed electrodes, the use of small amounts of sample, the high sensitivity and specificity, low limits of detection and the integration of nano materials that make it possible to improve the detected signal.

## 1. Introduction

In recent years, biosensors have emerged as an alternative to conventional methods due to their various advantages, such as low cost, portability, low detection limits, stability, the possibility of incorporating nanomaterials into their structure to amplify their electrochemical signal, and increased sensitivity and specificity. For these reasons, the development of sensors and biosensors had a significant peak, and their applications are increasing, in all industrial, environmental, biomedical sectors, etc. [1].

One of the fundamental parts of a biosensor is undoubtedly the electrode used as a support for mounting the system. Currently, much research is focused on the application of materials that improve the mechanical, electrical, and thermal properties of these electrodes. There is a variety of electrodes used as support in biosensor production, such as carbon paste, glassy carbon, gold, and screen-printed electrodes, with the latter standing out due to their advantages in working with micro volumes of samples and adaptability to portable devices. [1]–[4]

Composites have been widely used in the field of electrochemical sensor production, as such materials can provide improved chemical, physical, and magnetic properties, resulting in increased efficiency and sensitivity of detection systems [5], [6]. Among these composites, carboxylated graphene stands out among the class of chemically modified graphenes, as it presents graphite structures that are composed of a layer of graphene functionalized with groups such as hydroxyls, epoxides, carbonyls, and carboxylic groups. Its use for the functionalization of electrodes is based on some advantages they offer, such as good optical properties, large surface-to-volume ratio, excellent charge mobility and good electrical and thermal properties when compared to other carbon allotropes. Some advantages of using carboxylated graphene in the development of biosensors are based on its conductive nature, high surface area, and the presence of carboxylic groups in its structure, which can serve as anchoring points for biomolecules. [7], [8]

The coronavirus comprises a group of positive-sense single-stranded RNA viral material capable of infecting humans and animals, responsible for causing respiratory, gastrointestinal, neurological, and hepatic diseases that can range from mild symptoms to acute and chronic complications [9]. In the year 2019, cases of pneumonia of unknown origin were reported in the Chinese province of Wuhan. Subsequently, these cases were linked to the emergence of a new coronavirus, SARS-CoV-2, belonging to the group of betacoronaviruses [9].

With the advancement of the virus, the need for rapid and accurate diagnostic tests has become evident [10]. For the detection of the virus causing COVID-19, one can highlight tests such as RT-PCR (Reverse Transcription Polymerase Chain Reaction), considered the gold standard for SARS-CoV-2 detection. It is performed in the early stages of the disease due to the high viral load present in the respiratory tract during this period. Another factor is the need for specialized laboratories and personnel, in addition to its high cost. Consequently, alternative methods to the tests available on the market emerge as a solution, considering the limitations and high cost of tests like RT-PCR. Diagnoses through Point of Care Testing (POCT) can help alleviate pressure on laboratories and healthcare facilities, reducing the time to obtain a diagnosis, in addition to being cost-effective [11].

In this paper, carboxylated graphene was synthesized using a simple method and its morphological, spectroscopic, and electrochemical characteristics were studied. In addition, the obtained product was applied in the manufacture of a biosensor for the detection of IgA-SARS-CoV-2.

## 2. Materials and Methods

### 2.1. Reagents

Nitric acid (H_2_NO_3_), sulfuric acid (H_2_SO_4_), monosodium dihydrogen phosphate (NaH_2_PO_4_), disodium hydrogen phosphate (Na_2_HPO_4_), potassium chloride, N-hydroxysuccinimide ester (NHS), 1-ethyl-3-[3-(dimethylamino)propyl]-carbodiimide (EDC), and bovine serum albumin (BSA) were acquired from Sigma-Aldrich and Merck (USA). Graphene nanoplatelets was bought of XG sciences (USA). 1-Step™ Ultra TMB-ELISA substrate was purchased from Thermo Fisher™ Scientific (USA). N-protein was synthetized and purified in Federal University of Amazonas [12]. Also, 5 mmol L^-1^ K_3_Fe(CN)_6_/K_4_Fe(CN)_6_ prepared in 0.1 mol L^-1^ KCl was used as redox probe in the electrochemical measurements. All the electrochemical experiments were performed at room temperature (22 ± 0.5 °C) and without stirring. The human blood serum samples were provided by the Federal University of Amazonas, and this study was approved by The Research Ethics Committee of Federal University of Amazonas (CAAE:34906920.4.0000.5020) in accordance with Brazilian law and the Declaration of Helsinki.

### 2.2. Apparatus and Procedures

The carboxylated graphene was characterized using thermal gravimetric analysis (TGA) (Shimadzu TGA-50H, nitrogen atmosphere, flow rate of 50 mL/min, platinum cell, at 10 °C/ min). Scanning electron microscopy (SEM) (Tescan, model Vega 3) coupled to an energy dispersive X-ray spectroscopy system (EDS) (INCA energy, United Kingdom) operated with an accelerating voltage of 15 KV. The analyses by X-ray diffraction (XRD) were performed by Miniflex II Shimadzu, with with Cu kα radiation source, CuKα; wavelength, 0.15406 nm; diffraction angle, from 2° to 90°; the scanning step, 0.02°/min radiation source of 0.15406 Å, set at 2θ range 2–90°, with 2° min^-1^ scan speed, and Raman spectra was collected in a Horiba Scientific MacroRam spectroscopy equipment with output laser at 785 nm.

The electrochemical measurements were performed in an Autolab PGSTAT204 potentiostat/galvanostat equipped with FRA32M EIS module and NOVA software, 2.1.4 version (Metrohm Autolab, Netherlands). The commercial disposable SPCEs (ref. 110) from Metrohm Dropsens (Netherlands) was used as conductive support. They are composed of three electrodes, working electrode and counter electrode of carbon and reference pseudo-electrode of Ag. The electrochemical response was followed by cyclic voltammetry (CV) and electrochemical impedance spectroscopy (EIS), using [Fe(CN)_6_]^3-/4-^ as a redox probe to characterize the electrode surface after each modification with CG. CV was performed in a potential range between − 0.6 and 0.6 V, at scan rate of 50 mV s^−1^. The EIS measurements were executed in frequency range of 10 kHz to 0.1 Hz at potential of 0.15 V and amplitude of 5 mV.

### 2.3. Synthesis of Carboxylated Graphene

Firstly, we prepared the CG through a carboxylation method in acid medium (**Erro! A origem da referência não foi encontrada.**). For that, 7 mg of graphene was added to 30 mL of H_2_SO_4_(conc), and 3 mg was added to 10 mL of HNO_3_(conc). In the end, both solutions were mixed and sonicated for 1 h. Afterwards, it was made a centrifugation for 1 h at 22 000 rpm, in order to wash five times, the modified graphene with Milli-Q water. The main goal of this process was to neutralize the resulting graphene, until to achieve the pH 7 [13], [14]. The CG synthesized was then dried, using a rotary evaporation system.

Prior to modification, the carbon surface was electrochemically activated with 0.5 mol L^-1^ H_2_SO_4_ by CV technique, applying several potentials between −1.5 and 1 V at scan rate of 100 mV s^−1^ for five cycles [15], [16]. For the preparation of the CG/SPCE, the working electrode (WE) surface of the SPCE was modified with layers of CG, being deposited by drop-casting, 5 µL of a CG solution (2 mg mL^-1^) and dry at 60 °C.

### 2.4. Fabrication of the biosensor using CG

The biosensor for detect IgA of SARS-CoV-2 was fabricated according to the methodology described by Freire et al, [17]. The SPCE were modified with 3 layers of CG. Later, N-protein of SARS-CoV-2 was immobilized through the 1-ethyl-3-[3-(dimethylamino)propyl]-carbodiimide (EDC) and N-hydroxysuccinimide ester (NHS) coupling agent. This system allowed the detection of SARS-CoV-2 antibodies, IgA type, in human serum samples. The detection of the antibodies was performed using a reaction like enzyme-linked immunosorbent assay (ELISA) method. 7µL of human serum dilution (1:400 v/v) containing IgA-SARS-CoV-2 were deposited in the WE and kept in a humid chamber for 30 minutes. Afterwards, the immunosensor was incubated in horseradish peroxidase-labeled anti-IgA (1:300 v/v) for 20 minutes. After each modification step, the electrode was gently washed with PBS and Milli-Q water. For the enzymatic reaction was use 3,3’,5,5’-tetramethylbenzidine (TMB), being covered the area of the three electrodes (working, reference and auxiliar electrode) with 100 µL of TMB and kept in the dark for 2 minutes. The electrochemical response was followed by chronoamperometric (CA) technique at −0.19 V for 50s. The scheme 2 shows the steps described earlier.

**Scheme 1.**
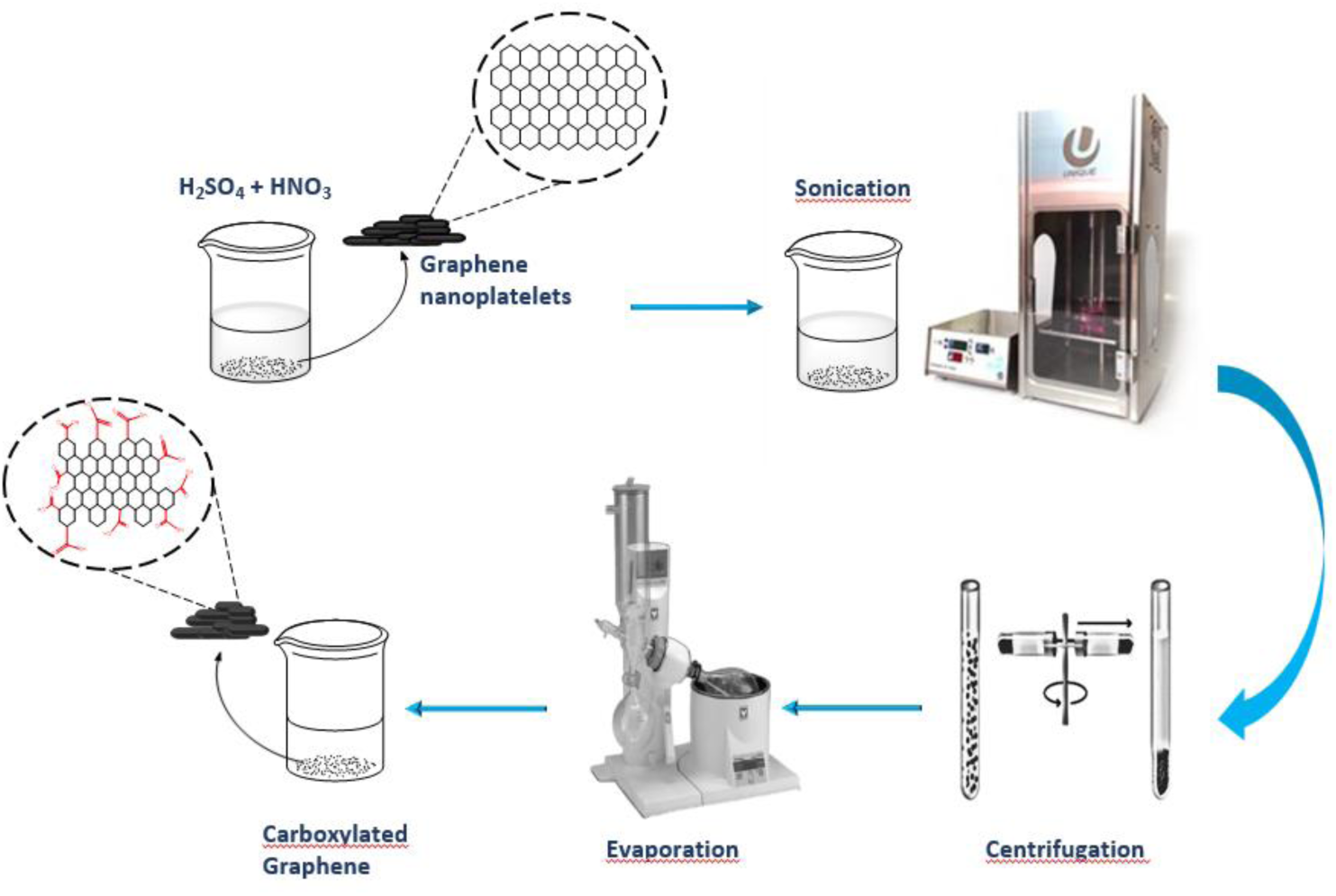
Representation of the method of synthesis of carboxylated graphene.

**Scheme 2.**
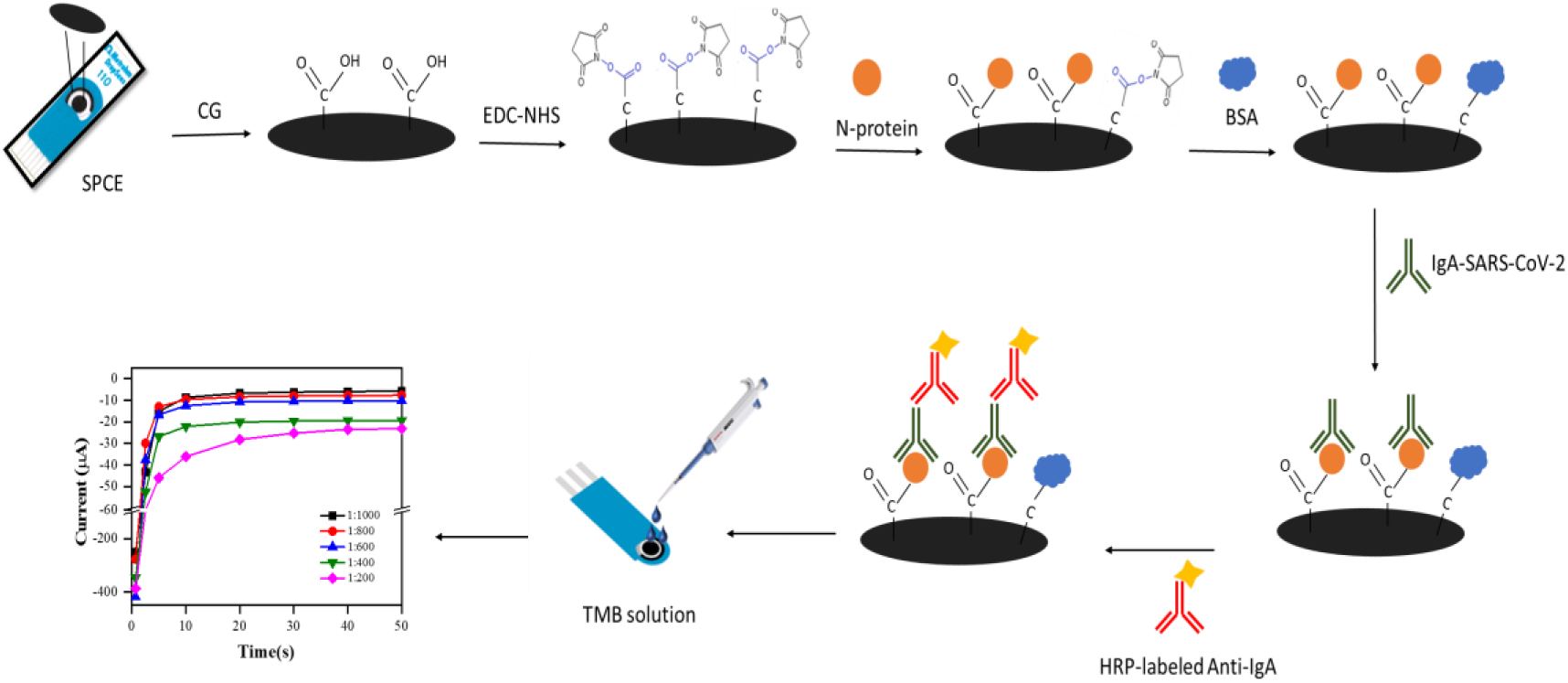
Schematic representation of the biosensor fabrication steps for IgA-SARS-CoV-2 determination.

### 3. Results and Discussion

#### 3.1.1. Characterization of the Materials

##### 3.1.1.1. **TGA**

Figure 1 shows the comparative TGA of graphene and carboxylated graphene. The decomposition was observed in three main steps: (I) evaporation of solvent and water, (II) decomposition of oxygenated functional groups (III) formation of CO. In the solvent evaporation step, which occurs between 0 °C and 170 °C, there is a mass loss of ∼ 3 % and 14 % for graphene and carboxylated graphene, respectively. These mass losses are related with the evaporation of solvent and constitutional water, and adsorption of gases on the surface of the material during the experiment [18]. The second stage, recorded between 170 °C and 610 °C, shows a greater mass loss (48%) for carboxylated graphene. In this temperature range, the decomposition of organic materials into carbonates occurs. The carbonization of organic cations can be considered as the main competitive reaction during the decomposition of organic matter, due to the oxidation of organic compounds in the air. In the last step, there is a small weight loss in the temperature range of 610 °C to 800 °C, approximately 1 % for graphene and 1 4% for carboxylated graphene, which may be associated with the decomposition of CO species [19].

**Figure 1.**
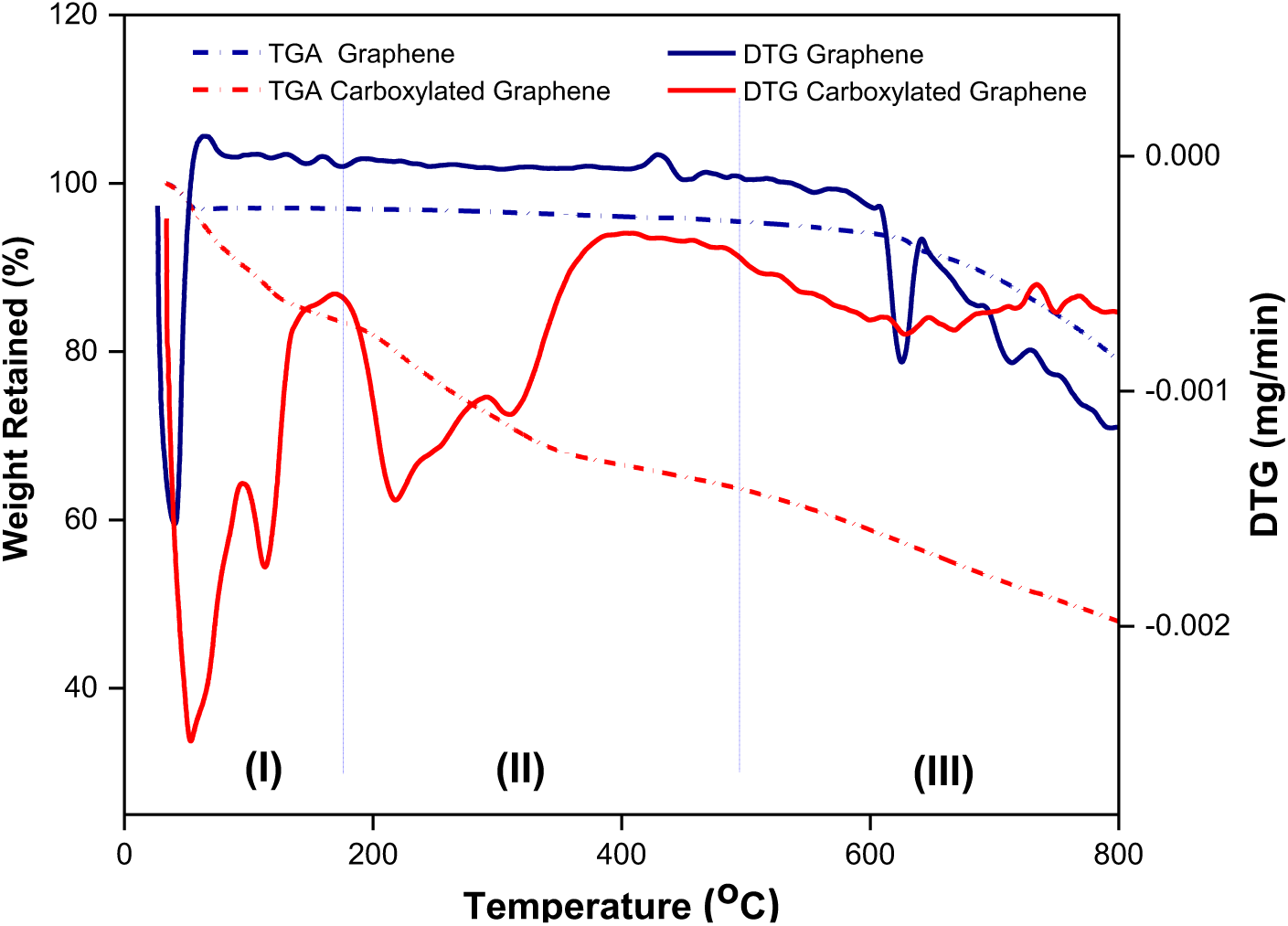
TG and DTG curves of decomposition of graphene and carboxylated graphene.

#### 3.1.2. X-ray diffraction

The X-ray diffraction spectra for graphene and carboxylated graphene are shown in Figure 2 (a) and(b).For spectrum 2 (a) it is possible to observe the presence of a more intense peak at 2θ = 11.64° corresponding to the presence of graphene oxide at an interplanar spacing of 7.8 x 10^-3^ nm. A wider peak is observed at 2θ = 24.35°, indicating the process of reduction of graphene oxide to graphene [20]. For the spectrum represented in Figure 2(b) it is possible to observe the peak 2θ = 11.80° corresponding to graphene oxide and it is possible to observe a peak at 2θ = 26.44° indicating the increase in crystallinity in relation to the observed process that can be justified due to the presence of carboxylic groups in the material.

**Figure 2.**
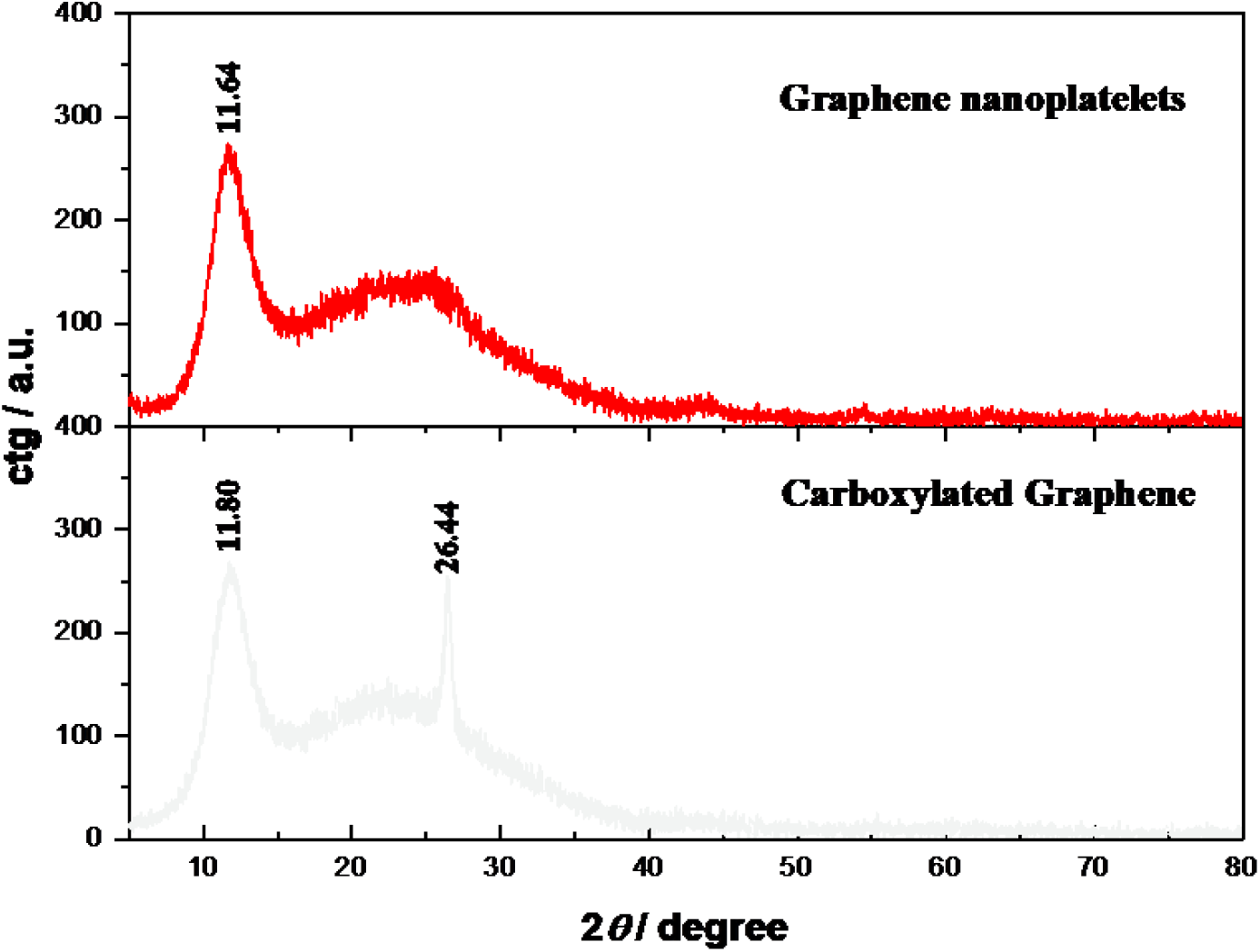
X-ray diffraction pattern of graphene and carboxylated graphene.

#### 3.1.3. Raman spectroscopy

Raman spectra shown in Figure 3, (a) graphene nanoplatelets and (b) carboxylated graphene were obtained at room temperature. Figure 3a shows the three peaks located at 1065, 1345 and 1598 cm^-1^ in this material is supported by the characteristic D*, D, and G modes, respectively [21]. In figure 3b, the three characteristic bands of carbon-based materials, which correspond to the band D, G, and 2D bands centered at ∼1320, 1585 and 2640 cm^-1^, respectively [5]. The band referred as D* is related to disordered graphitic lattices provided by sp^2^-sp^3^ bonds [21]; The G band, it represents the planar configuration sp^2^ bonded carbon that constitutes graphene; The D band is known as the defect band or the disorder band. The band is the result of a one phonon lattice vibrational process. At 2D band is corresponded to a two phonon lattice vibrational process, similar materials such as carbon nanotubes [21]–[23].

**Figure 3.**
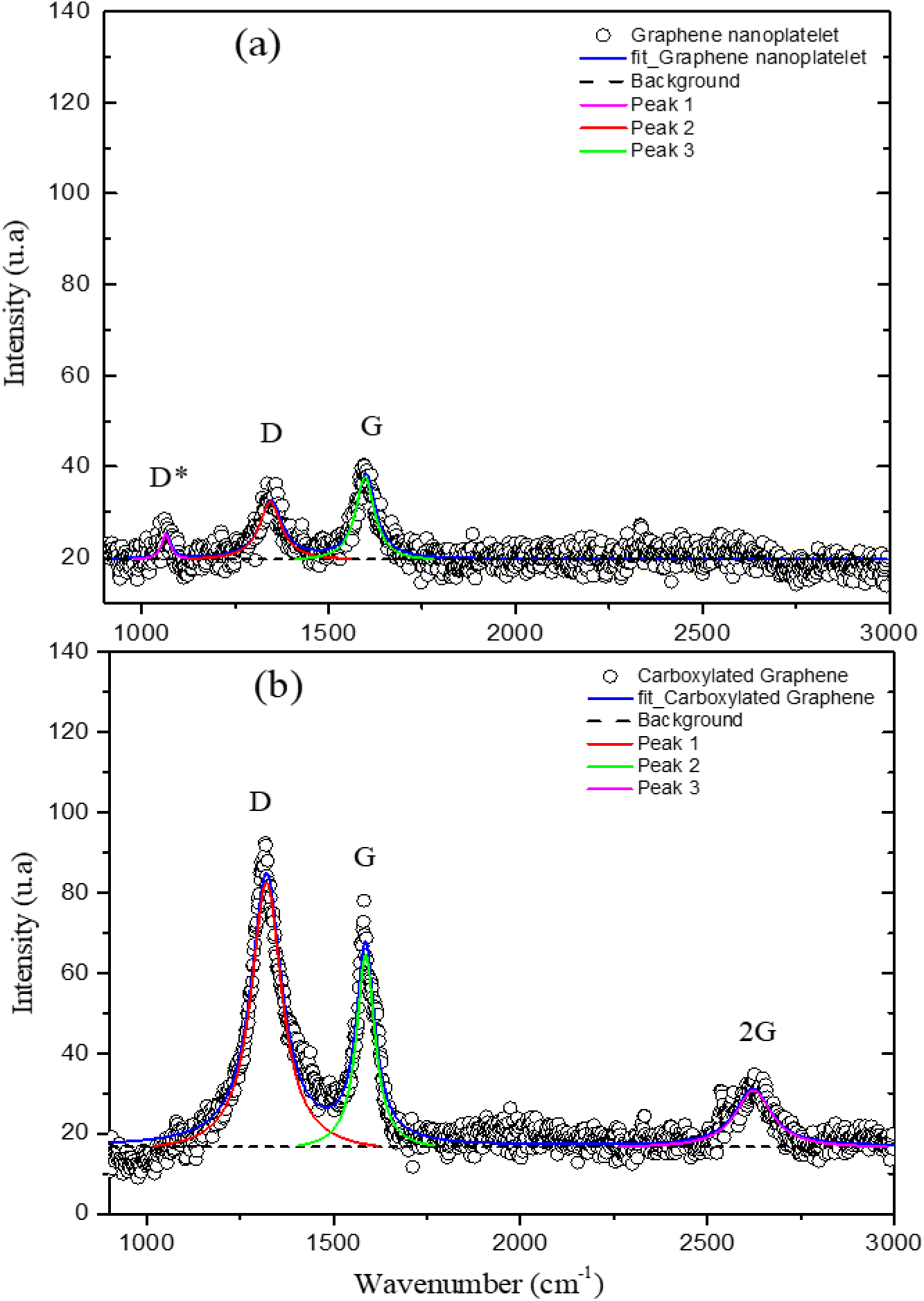
Raman spectra of (a) graphene nanoplatelets and (b) carboxylated graphene with the D*, D, G and 2G band deconvolution.

The intensity ratio of the D band to G band (ID/IG) characterize the disorder degree of graphene, and for instance for graphene nanoplatelets at 0,7 and carboxylated graphene at 1,3, as shown in Table 1. The ratio of ID’/ID is used to explore the types of product defects generated, for graphene is 1,3 indicating defect on bounds and for carboxylated graphene indicating vacancies in the hexagonal structure [24]. This result is in according with the observed by XRD pattern, the process of graphene carboxylation increases of amorphism.

**Table 1.** Details of Raman spectra of nanoplatelets and carboxylated graphene. The peak intensity ratio of ID/IG.

| Samples | D peak<br>(cm<sup>-1</sup>) | $\beta$<br>(cm<sup>-1</sup>) | G peak<br>(cm<sup>-1</sup>) | $\beta$<br>(cm<sup>-1</sup>) | I<sub>D</sub>/I<sub>G</sub> |
| --- | --- | --- | --- | --- | --- |
| Graphene nanoplatelets | 1345 | 12.8 | 1598 | 18.5 | 0.7 |
| Carboxylated graphene | 1322 | 67.6 | 1586 | 49.0 | 1.3 |

**Table 2.** Rct values calculated for the different depositions using the Randles equivalent circuit.

| Number of Depositions | Rct ($\Omega$) |
| --- | --- |
| Bare SPCE | 240 |
| 1 | 81 |
| 2 | 71 |
| 3 | 44 |
| 4 | 33 |
| 5 | 22 |
| 6 | 19 |

#### 3.1.4. SEM and EDS

The SEM images of graphene and carboxylated graphene are shown in **Erro! A origem da referência não foi encontrada.**. The imagens (a-b) reveals that the graphene powder had a flake structure of non-uniform size with a slightly rough surface and numerous obvious folds. On the contrary, the CG (Figure 4 c-d) has greater uniformity in relation to graphene, the general structure of the CG is quite altered, it is verified that the thin plates are well exfoliated, with a wrinkled and crushed appearance of the leaves that was evident. The prepared CG surface was smoother, more transparent, and thinner than graphene. The results showed that the CG was successfully prepared by reaction in an acid medium [24], [25].

**Figure 4.**
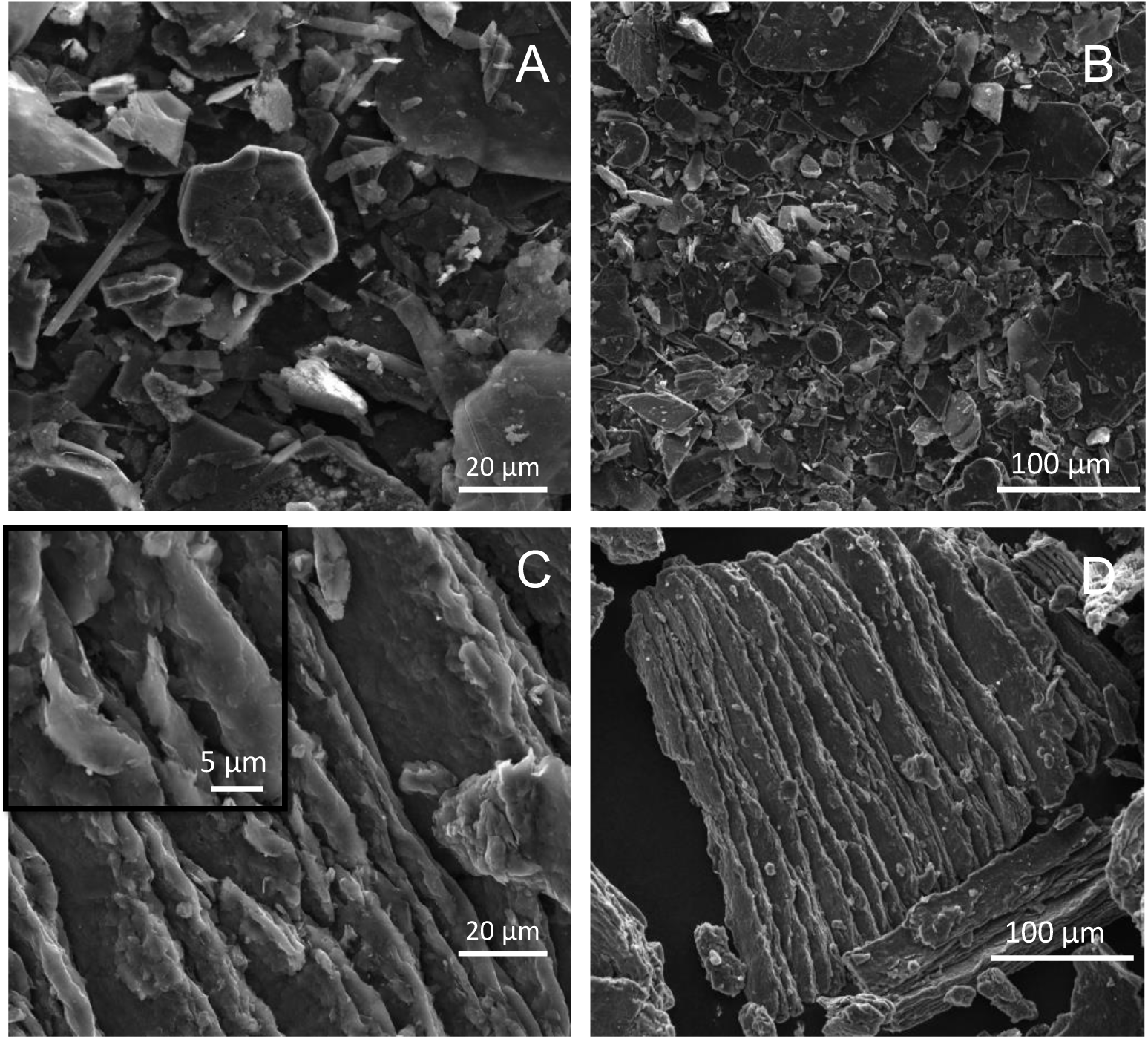
The SEM images of a graphene (A-B) and a carboxylated graphene (C-D).

To verify the success of the synthesis, energy dispersive spectroscopy coupled with scanning electron microscopy (SEM-EDS) was used (Figure 5). The presence of only element C in the graphene spectrum, indicating high purity of this sample. The elemental composition demonstrated in the characteristic peaks of C and O present in CG the presence of carboxylic moieties available for further binding.

**Figure 5.**
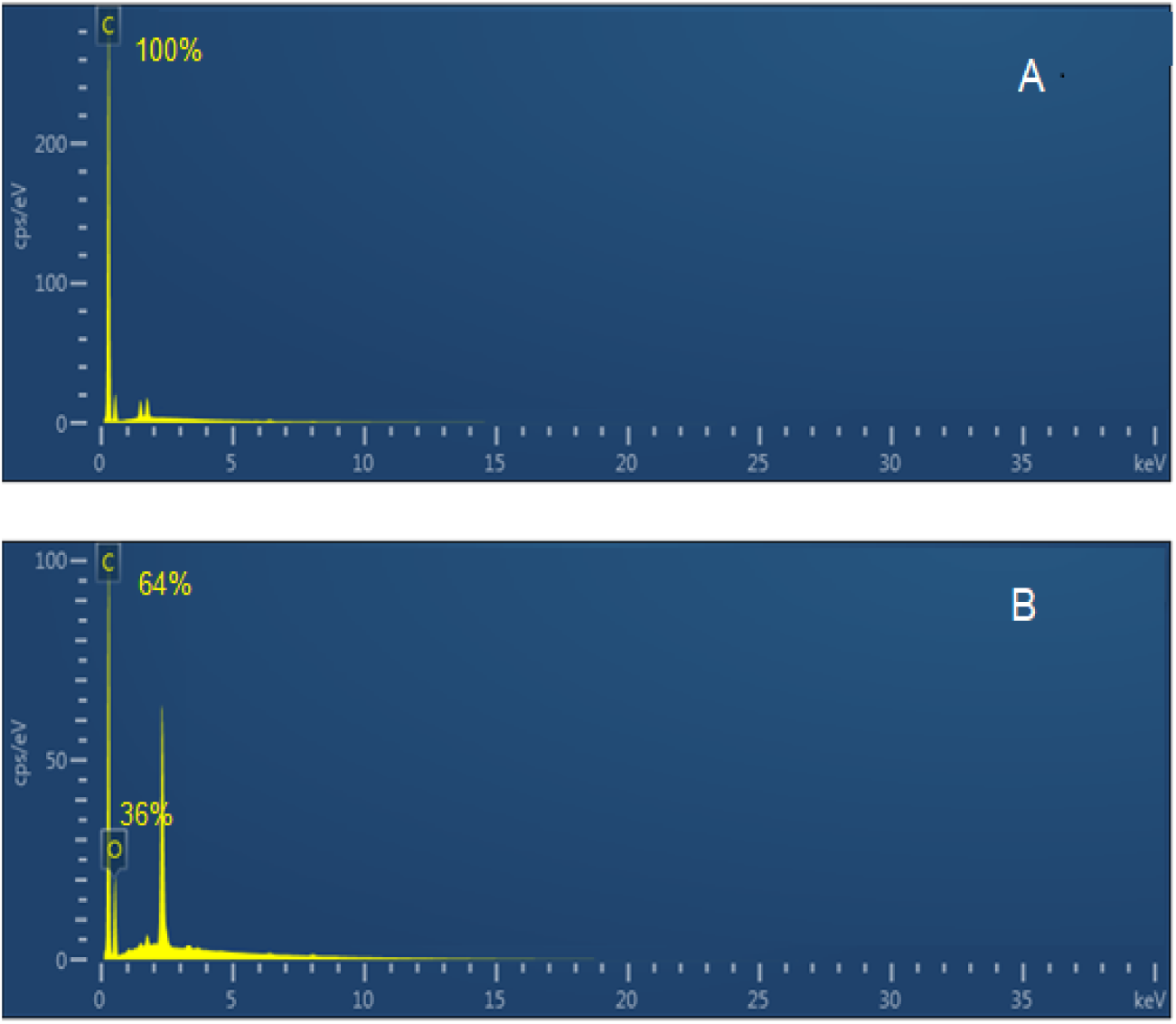
EDS spectrum of the (A) graphene and (B) carboxylated graphene.

### 3.2. Electrochemical properties

Chemically customized electrodes containing surfactant species showed prominent advantages over bare electrodes. It is feasible to modify commercially available electrodes to provide good physical distribution, increased electrochemical signal and immobilized surface. To determine the ideal number of depositions to obtain an improvement in electrochemical signals, 6 depositions were examined, each with a volume of 5µL of the distribution of 2 mgmL^-1^ of CG on the SPCE surface. The electrochemical behavior of electrodes modified with CG classes was investigated using the CV technique (Figure 6 A).

**Figure 6.**
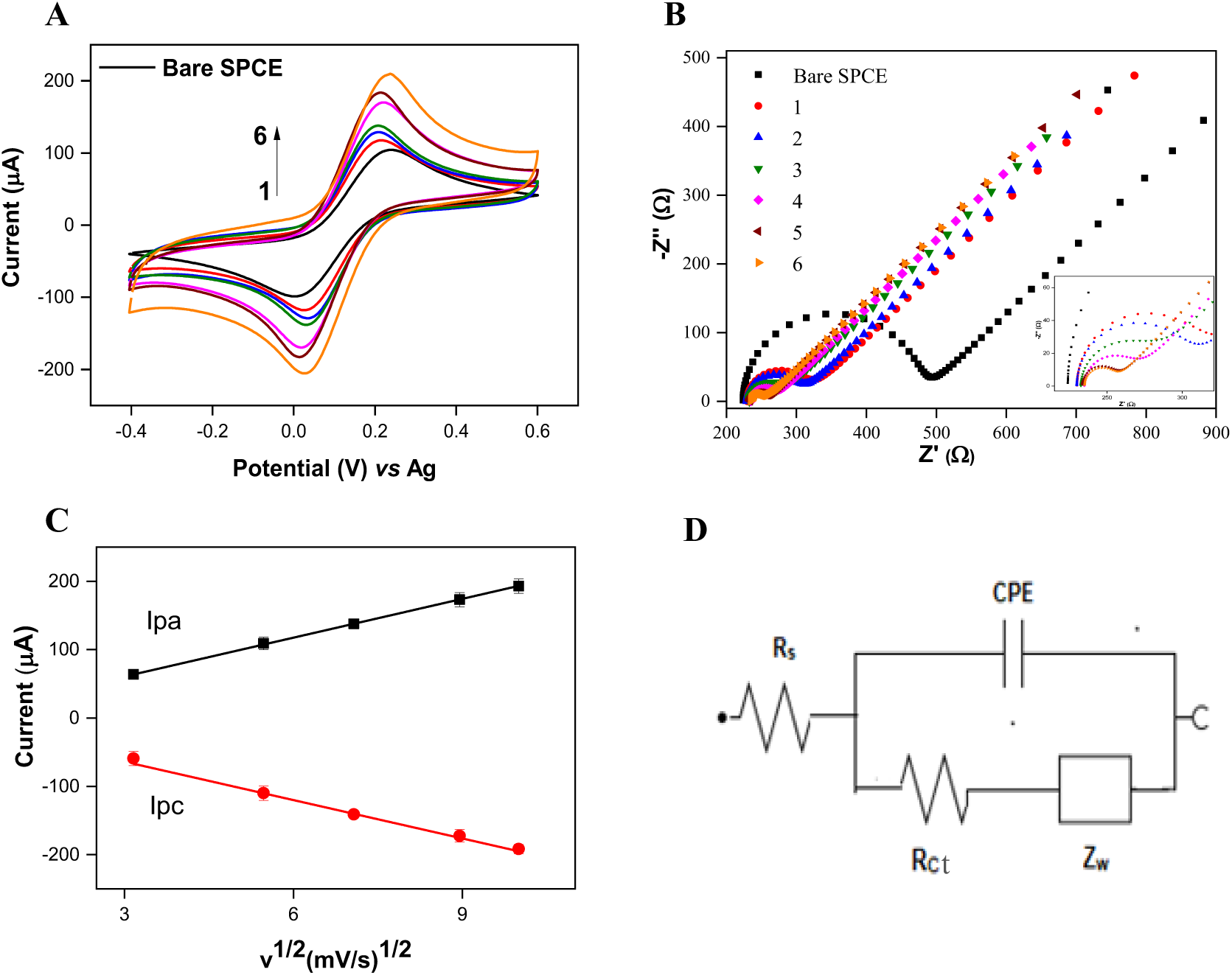
(A and B) CV and EIS for deposition of carboxylated graphene layers in 5 mmol L^-1^ [Fe(CN)_6_]^3−/4−^ + 0.1 mol L^-1^ KCl. I Linear relationship between current and square root of sweep speed (10 – 100 mV s^-1^) for modified SPCE with 3 depositions of CG. (D) Randles equivalent circuit used in EIS.

As observed, the electrochemical signal increases with each added layer of CG. With increasing current, it is observed that the presence of carboxylated graphene significantly improves the intensity voltametric signals due to the properties of these materials, especially their high electrical conductivity and electroactive surface area [26], [27]

In the case of bare SPCE, the redox peaks exhibited ∼0.236 V of separation. After the CG modification layers, the peak currents of CG/SPCE increased between 12-102%. Furthermore, the anodic and cathodic peak potentials approached each other, showing a decrease in ΔEp of about 0.–6 - 0.2 V, which suggests an increase in electron transfer between the electrochemical probe and the surface of the modified electrode. Consequently, the electrochemical performance of SPCE was improved.

When modifying the electrode with CG, there was an increase in the capacitive current directly related to the increase in the electroactive area and the roughness factor. The capacitive current is generated due to the presence of an accumulation of electrons on the surface of the electrode, increasing the charge of the electrical double layer. Graphene is one of the most used carbon nanomaterials due to its high electrical conductivity and large surface area compared to carbon nanotubes and graphite. Although graphene can be used to improve the electrical conductivity of composite materials, it also causes very high capacitive currents from SPEs (> 1 µA). To minimize these effects, a potential must be fixed during electrochemical measurements, and the capacitive current contribution is minimized, therefore, the analytical detection in biosensors suggested for SPCE modified with this material is chronoamperometry [28], [29]

The EIS analysis (Figure 6 B) showed changes in resistance charge transfer represented in the Nyquist plot. The Nyquist diagrams can be simulated with an equivalent circuit (Figure 6 D), where R_S_ is the resistance of the electrolyte solution, CPE is the constant phase element, Rct is the electron-transfer resistance, and Z_W_ is the Warburg element due to diffusion of the redox couple ([Fe(CN)_6_]^4-^/[Fe(CN)_6_]^3-^) to the interface from the bulk interface from the bulk of the electrolyte. The semicircle portion from high to intermediate frequencies refers to the kinetic charge transfer process whereas the straight line at low frequencies arises from the diffusional barrier regarding the redox couple mass-transfer [30].

The EIS spectra show clear differences in signal after integration of each layer onto the SPCE surface starting with the bare SPCE (Rct = 240 Ω), which exhibited a semicircle with a small diameter at high frequencies representative of some SPCE resistance to electron transfer. After modification with CG, there was a gradual decrease in resistance (show in **Erro! A origem da referência não foi encontrada.**2) to electron transfer confirmed by the disappearance of the semicircle which was replaced by a straight line indicating a diffusion process. This is due to the increase in conductivity, thus improving the transfer of electrons between the solution and the surface of the modified electrode.

The experiment using different scan rate were carried out with the objective of evaluating the reversibility and the nature of the transport involved in the SPCE type electrode modified with CG. The CV analysis allowed characterizing the surface of the SPCE modified with three layers of CG. The electrochemical behavior of the electrode evidenced a good performance, and low increase in ΔE when increasing scan rate, allowing the conclusion that the reaction is quasi-reversible. As can be seen in Figure 6 C, with increasing scan rate, the anode peak currents increase linearly according to the following equations:

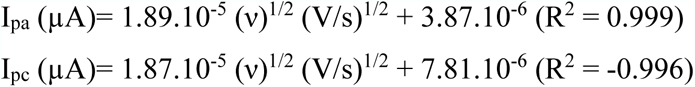

### 3.3. Electrochemical Immunosensor based in CG

The biosensor assembly steps were characterized by electrochemical techniques (CV and EIS), and the experimental variables were optimized to obtain the best device performance. These results were reported in the work published by Freire et al, [17].

#### 3.3.1. Detection of IgA-SARS-CoV-2

For the detection of IgA-SARS-CoV-2, human serum samples from negative and positive patients for Covid-19 were used. In the experiments, different concentrations of positive serum were tested, varying the dilution factor from 1:1000 to 1:200 v/v in PBS 1X. The Figure 7 A shows the results of chronoamperometric measurements performed at -0.19 V (reduction potential) vs. pseudo-Ag for 50s. During the first 10s, a rapid decrease in current values was observed, due to electrode polarization. After 10s, the reaction stabilizes and begins to reach a plateau state. The chronoamperograms registered for the blank (PBS) and negative sample were similar, with current values close to zero, since there is no enzymatic reaction, a behavior different from that recorded by the positive sample, where there was an increase in reduction currents (Figure 7 B)

**Figure 7.**
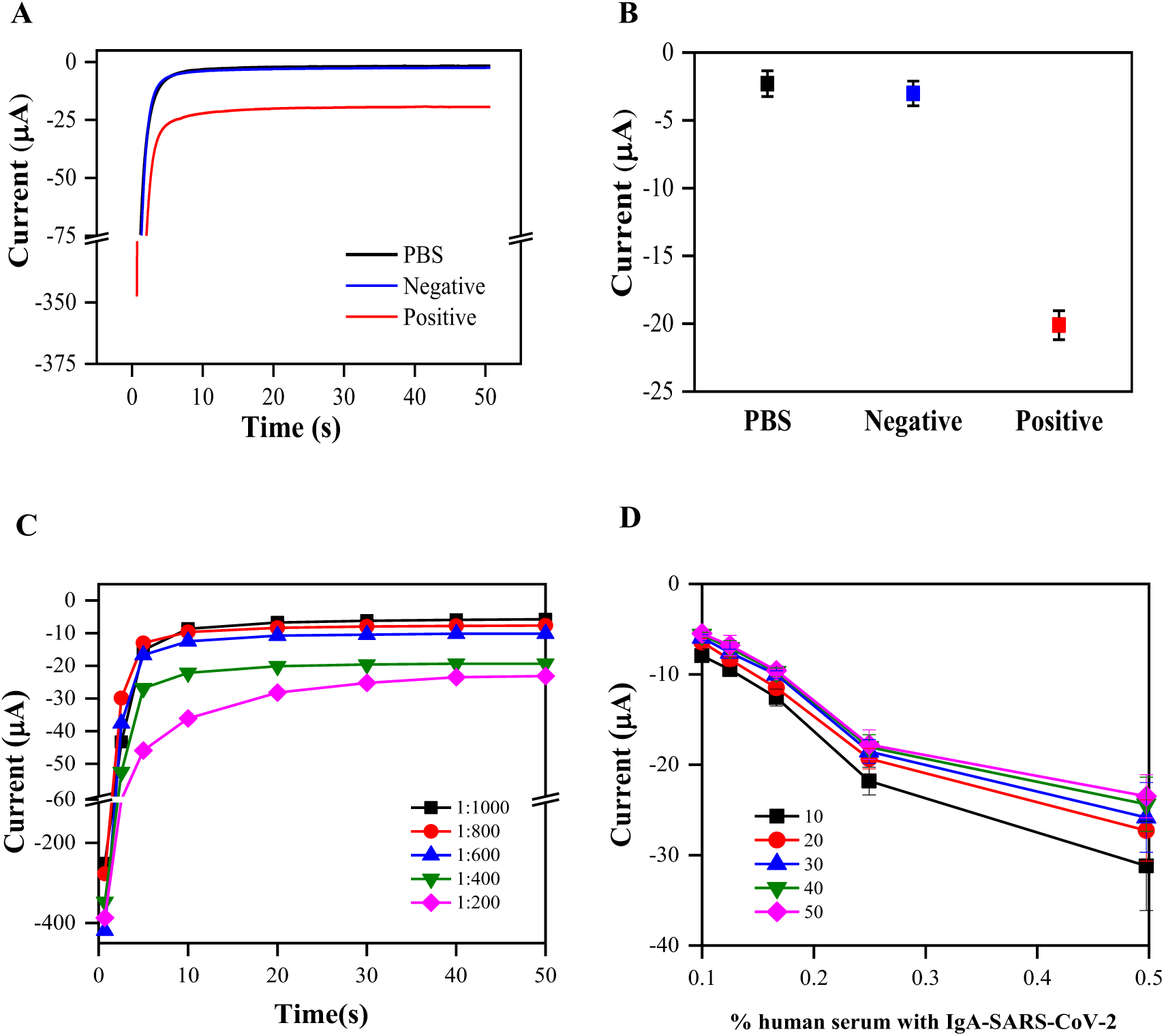
(A and B) Chronoamperograms recorded with PBS, negative and positive controls. (C) Chronoamperometric response measured for different concentrations of human serum with IgA-SARS-CoV-2. (D) Current values registered from 10 to 50 s for different concentrations of serum human with IgA-SARS-CoV-2.

The response of the positive sample at different concentration is showed in Figure 7 C, recording a linear relationship between % human serum with IgA-SARS-CoV-2 and the current values. The current was analyzed from 10 to 50s (Figure 7 D), being observed this linear behavior in all time periods. However, the linearity decreased at the highest IgA-SARS-CoV-2 concentration, what could be a consequence of the electrode passivation related to TMB precipitation [17], [31], [32].

The calibration curve was plotted using absolute values of stable current recorded at 20s against % human serum with IgG-SARS-CoV-2, and the performance was evaluated at 1:1000; 1:800; 1:600; 1:400; 1:200 v/v dilution factor. The regression equation obtained was y= 58.5038 x + 3.43654, with R^2^ = 0.94 (Figure 8 A). The limit of detection (LOD) was calculated according to the International Union of Pure and Applied Chemistry recommendations, where LOD = 3 x SD/m (SD: standard deviation blank and m: slope of the calibration curve)[33]. This limit was established as 4.59 µA, corresponding to a % IgG-SARS-CoV-2 of 0.0624, which is equivalent to a dilution factor of 1:1601 v/v.

**Figure 8.**
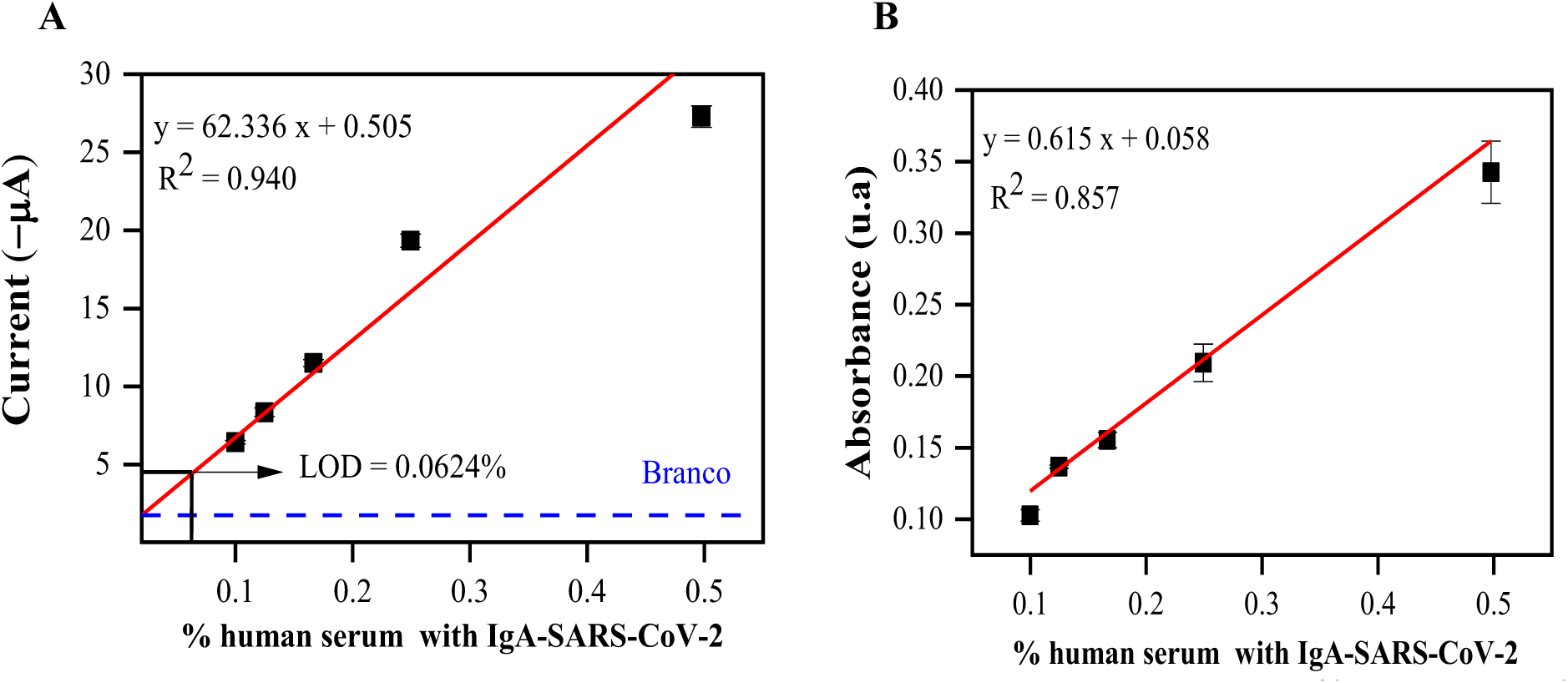
Calibration curve for IgA-SARS-CoV-2 determination through (A) CA technique and (B) ELISA technique. Standard error bars correspond to correspond to measurements made on three replicates of each concentration (n = 3).

Experiments using ELISA technique were carried out with the purpose of validating the method presented in this work. The calibration curve (Figure 8 B). was obtained by plotting of absorbance versus % serum human with IgG-SARS-CoV-2. The results exhibited a linear relationship in the same concentration range to our method (from 1:1000 to 1:200), with a LOD = 0,0771% of IgG-SARS-CoV-2, which is equivalent to a dilution factor of 1:1296 v/v. The curve equation was defined as y= 0.615 x + 0.058, with R^2^ = 0.857. It should be noted that the immunosensor recorded better values of R^2^, the LOD was 1.2 times greater, used half of the total time for application of the methodology and smaller amount of immobilized protein and volume sample when compared with the ELISA assay.

## 4. Conclusions

In this work, a simple, straightforward, and low-cost method of synthesis was used for obtaining CG. The product present good conductivity, which is ideal for application in the development of electrochemical biosensors. A platform using immobilization of biomolecules (N-protein of SARS-CoV-2) onto CG/SPCE was constructed for detecting of IgA antibodies, that are marker of the covid-19 disease. The device responded sensitively and selectively to the target analyte, and the results of our method were confirmed by ELISA test, which is a standard method.

